# A covariant model of molecular and diversification patterns through time and the history of large clades

**DOI:** 10.1101/2024.02.01.578373

**Authors:** Graham E. Budd, Richard P. Mann

## Abstract

Rate shifts in speciation and extinction have been recognised as important contributors to the creation of evolutionary patterns. In particular, the distribution of modern clade sizes is difficult to reconcile with models that do not include them. Although recent advances have allowed rate shifts to be integrated into evolutionary models, these have largely been for the purpose of inferring historical rate shifts across phylogenetic trees. In addition, these models have typically assumed an independence between patterns of diversification and rates of molecular and morphological evolution, despite there being mounting evidence of a connection between them. Here, we develop a new model with two principal goals: first, to explore the general patterns of diversification implied by constantly changing rates, and secondly to integrate diversification, molecular and morphological evolution into a single coherent framework. We thus develop and analyse a covariant birth-death process in which rates of all evolutionary processes (i.e. speciation, extinction and molecular and morphological change) covary continuously, both for each species and through time. We use this model to show that modern diversity is likely to be dominated by a small number of extremely large clades at any historical epoch; that these large clades are expected to be characterised by explosive early radiations accompanied by elevated rates of molecular evolution; and that extant organisms are likely to have evolved from species with unusually fast evolutionary rates. In addition, we show that under such a model, the amount of molecular change along a particular lineage is essentially independent of its height, which further weakens the molecular clock hypothesis. Finally, our model predicts the existence of “living fossil” sister groups to large clades that are both species poor and have exhibited slow rates of morphological and molecular change. Although our model is highly stochastic, it includes no special evolutionary moments or epochs. Our results thus demonstrate that the observed historical patterns of evolution can be modelled without invoking special evolutionary mechanisms or innovations that are unique to specific times or taxa, even when they are highly non-uniform: instead they could emerge from a process that is fundamentally homogeneous throughout time.

## Introduction

Contemporary modelling of patterns of evolutionary history considers the overall process that generates it to consist of three major components: the nature of molecular substitution; its rate of change; and the nature of the branching process itself [1]. Notably, each of these three components are typically treated as being independent, so that, for example, the rate of branching is not correlated with the rate of molecular change (e.g. [2]). Nevertheless, there are sound empirical and conceptual reasons for thinking that at least the second and third of these may be intimately related [3], and attempts have sometimes been made to consider them jointly (e.g. [4] [5]). Although various models for the branching process exist, one of the most commonly used is the birth death process (BDP) [6]. One obvious discrepancy of real data compared to basic BDPs is that diversification rate shifts must surely have occurred repeatedly through time, as a single homogeneous BDP cannot possibly capture the true patterns of diversification reflected in evolutionary history (c.f. [7]). Rate shifts have largely been addressed in one of two ways: either by assuming rate shifts occur at significant rare points (the “key innovation” concept), or by assuming broad secular variation, e.g. with declining rates through time across the entire tree [8] [9]. A recent paper by Maliet et al. [10] took a more innovative approach by developing a model of frequent small rate shifts associated with speciation events through time (for other non-continuous models, see [11] and the review in the supplementary information of [10]). Their model has the advantage of negating the need of identifying the key moments when such shifts took place, and instead takes a more satisfying homogeneous view of rate shifts through time (c.f. [12]). They neatly explained why observed clades are often imbalanced and “stemmy”, but were otherwise focused on developing a framework of *inference* of rates through time within such a model, based on molecular data from extant taxa [13]. Our aim here is to develop a more general *forward* model of diversification with small rates shifts, in order to explain more broadly the observed characteristics of extant clades that are not well captured by traditional BDPs, and to then tie them into potential patterns of molecular evolution. In particular, two interesting features of evolutionary patterns stand out as being in need of explanation: i) the distribution of clade sizes, and ii) variation of molecular rates of evolution along the lineages, especially at the beginning of clades. Our purpose here is thus not to infer historical patterns associated with particular trees, but to reveal what the large-scale patterns of evolution associated with continuous small rate shifts through time are.

The distribution of modern diversity predicted by BDPs (homogeneous or epochally time-varying across all species) is geometric [8] [14], and this remains the case even when non-selective mass extinctions are considered [15]. However, a certain amount of evidence suggests that extant sizes are in fact over-dispersed relative to this expectation. Consider, for example, the crown group animal phyla, which for the sake of argument we can assume all emerged around 500 Ma. Estimating total species diversity in the phyla is fraught with difficulty, but even so the species count differs widely. For example, the phyla have an average diversity of c. 50,000 species, but the arthropods have a diversity of well over one million species, thus being over twenty times larger than expected. Under a geometric distribution this is essentially impossible (*p* ∼10^*−*7^). This pattern is seen repeated hierarchically: e.g. most arthropods are insects, and most insects appear to be hymenopterans [16]. Similarly, the angiosperms are much more diverse than any other plant clades (e.g. c. 300000 versus 1000 gymnosperms) and birds much more so than crocodiles in the archosaurs (c.10000 versus c. 85). In other words, the existence of Stanley’s “supertaxa” [17] does not seem compatible with a purely geometric distribution of clade sizes as predicted by the homogeneous BDP. In addition, clade sizes show a complex relationship with age that is not easily explained by homogeneous diversification (e.g. [18][19][20]), and indeed attempts to estimate absolute rates within a clade suggest several orders of magnitude variation [19]. Although there been some argument about this in the literature then, it seems that clade sizes do often appear overdispersed relative to any expected geometric distribution.

A second anomaly is the problem of non-homogeneity in rates along the lineages (i.e. those historical species that in retrospect led to at least one modern survivor). At least some studies have suggested the “early burst” model of evolution, i.e. that rates of molecular (and sometimes morphological) evolution are high in early lineages, and quickly decline after (see e.g. [21][22]). Examining early burst evolution empirically is complicated by elevated early molecular evolution rates making clades look older than they actually are [23][11]). A prominent study that indicated such early elevated rates was that of Lee et al. 2013 on arthropods [21]. When they fixed the age of the arthropod clade to conform with the fossil record, elevated rates of both molecular and morphological evolution were observed in the early lineages. One approach to resolve this problem, which is one they themselves investigated, is to break the link between the fossil record and the age of the clade. In such a case, the apparent elevated early rates can be explained by “bunching up” the early lineages to artificially squeeze the clade into a too-narrow time interval (c.f. [24]). However, we have previously marshalled strong reasons for thinking that the fossil record in such instances is often reliable, in which case early bursts should be taken seriously and not dismissed as dating artefacts [15][25][26]. It has been previously shown that so-called “large clades” (i.e. clades that produce more species than expected in the present day, given their diversification parameters) are expected to exhibit early rapid bursts of at least *lineage* creation [27][28]. However, this results from greater-than-expected long-term survival of new clades, and not from substantially increased diversification rates along the lineages (which might be considered to be linked to increased rates of substitution). In such clades, there must be at some point higher rates of diversification overall, and this is most likely to be at the start; but this increased rate is accommodated by more lineages surviving, and not by markedly increased rates of diversification along those lineages. The upshot of this result is that under a standard BDP, one would not necessarily expect markedly increased rates of molecular (or morphological) evolution in early lineages, even in unusually large clades. Any correlation between rates of evolution and the BDP is unlikely to be merely coincidental, although it has been noted before that formally considering the two processes together is rare in the literature [29][4]. A considerable body of evidence now exists that does indeed suggest such a link between diversification and molecular rates of evolution (e.g. [30][29][3]) and have proposed several ways in which this link might be created. We will thus also consider how the model we describe below might be expected to influence rates of molecular (or morphological [31]) evolution through time.

## Model outline

The problems highlighted above suggest that neither a homogeneous BDP nor one with epochal change across all species, are sufficient to explain some important empirical features of diversification. To address these problems we have created a BDP model in which rates of speciation and extinction continuously co-vary in individual species, rather than across the whole tree as a function of time. Under this model, all evolutionary rates are specific to a given taxon at a specific moment in time. Our model can be seen as an extension and further development of an important recently-analysed model [10] (c.f. [5]) that proposed a BDP in which each species has its own diversification rate. Here, we propose that there exist baseline rates of speciation (*λ*_0_) and extinction (*µ*_0_) that are linearly modulated by a new variable we label as *tempo, τ*, which controls the relative rates of all evolutionary processes. At any given time a taxon with tempo *τ* has a speciation rate *λ* = *τλ*_0_ and an extinction rate *µ* = *τµ*_0_. In contrast to previous approaches that have considered rate shifts, we neither assume that shifts occur at ‘special’ moments in time [32] or that they are necessarily associated with speciation events [10]; instead the tempo varies continuously through time, and independently along all branches of the tree. In addition to arguably being more ‘natural’ in the sense that this removes all reference to particular events, this produces a mathematical structure that is highly tractable for answering questions about the dynamics of the process.

This model is fully covariant, in that all rates are linked directly to *τ* ; in effect the tempo represents a local speeding-up or slowing-down of evolutionary time, such that all processes happen faster or slower. In particular, we assume that tempo *itself* varies through time, and the evolution of *τ* itself proceeds at a rate proportional to *τ*. Specifically we model the log-tempo (*x* = log *τ*) as evolving according to a modified Ornstein–Uhlenbeck (OU) process that incorporates the effect of the tempo itself on all rates:

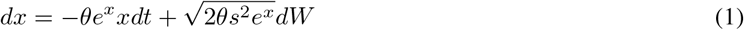

where *dW* represents an incremental change from a Wiener process. We impose this model for the evolution of the log-tempo *x* since the tempo itself is constrained to be positive. Symmetric evolution of *x* implies a scaling symmetry for the tempo, such that rates of evolution are equally likely to rise or fall by any fixed scaling factor. The parameters of this stochastic differential equation are the mean reversion rate *θ* and the stationary variance of the process, *s*^2^. The *e*^*x*^ terms in this equation come from the self-interaction of the tempo, which as well as multiplying the rate of all other processes also determines the rate at which it evolves itself, such that the effective increment of time is *e*^*x*^*dt*.

As we show in Methods, this results in a drift-diffusion partial differential equation for the generating function of the resulting birth-death process:

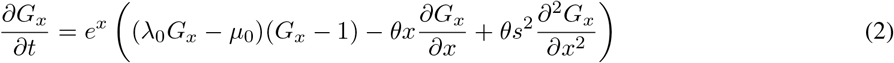

where 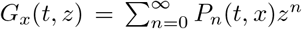, with *P*_*n*_(*t, x*) being the probability of generating *n* species over time *t* in a process starting with log-tempo *x*. Solving this equation for an initial condition *G*_*x*_(*t* = 0, *z*) = *z* provides the value of the generating function *G*_*x*_(*t, z*).

Equation 2 does not appear to permit solution in closed form, but can be straightforwardly solved numerically. The values of *P*_*n*_(*t, x*) can be retrieved from this generating function by Fourier inversion (see Methods).

We can derive further equations specifying the evolution of the mean number of species generated by the process over time, the expected number of lineages (species that will have modern descendants) and the distribution of tempos over time. Derivation of these equations is described in Methods. The most important of these equations specifies the evolution of the mean number of species through time. Given a generating function *G*_*x*_, the mean of the distribution, *N*_*x*_(*t*) ≡ 𝔼(*n* | *t, x*) is given by:

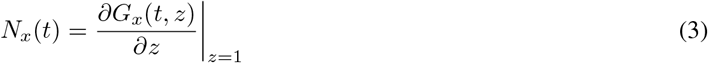

Using this relation, equation 2 can be transformed into a simpler, linear form to represent the dynamics of the mean:

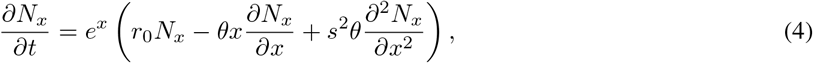

where *r*_0_ = *λ*_0_−*µ*_0_ is the net diversification rate. This equation reveals the key dynamics of the process: the expected number of species with log-tempo *x* locally increases exponentially at the rate *r*_0_ modulated by *τ* = *e*^*x*^. At the same a drift-diffusion process modifies the tempo of each species, such that species tend to move towards a log-tempo of 0 (i.e. *τ* = 1).

## Results

We analysed our model with core set of parameters *λ* = 0.51, *µ* = 0.5, *θ* = 0.01, *s* = 1. It is not our goal to quantitatively fit our model to the modern diversity or evolutionary history of any specific clade, but rather to reveal the qualitative features the model predicts. These parameters are chosen to reflect reasonable expectations about the real evolutionary process: a baseline extinction rate of *µ* = 0.5 comports with that chosen in previous analyses (e.g. [28]) and, combined with a speciation rate of *λ* = 0.51 is consistent with a typical species existing for c. 1 myrs, in broad agreement with the fossil record. The speciation rate is chosen to be of similar magnitude to the extinction rate, such that extinction plays a significant role in the evolutionary dynamics, but is otherwise arbitrary. We choose a mean-reversion parameter *θ* = 0.01 to be equal to the net diversification rate as we will later show that if *r*_0_ = *θ* then the mean log-tempo converges to 0 (see Methods, equation 37). Although this choice is mathematically convenient we do not expect it represents any necessary feature of the evolutionary process, nor do the qualitative features of our results depend on it. Finally the diffusion parameter *s* = 1 is chosen to be large enough produce significant effects of the diffusive dynamics, and otherwise is simply a mathematically convenient choice.

### Distribution of clade sizes

We solved equation 2 for times 0 ≤*t* ≤500 myrs and starting log-tempos −10 *< x <* 10 and performed a Fourier inversion (see Methods) to retrieve the implied probability distribution *P*_*n*_(*t* = 500myrs, *x*). The distribution of clade sizes for a clade that starts with log-tempo *x* = 0, excluding clades of size zero, is shown in Figure 1A. The clade sizes follow a distribution that differs strongly from the geometric distribution expected under a typical BDP. This distribution is characterised by most clades being small, but with a few extremely large clades. This means that clades that are many times greater than average (either mean or median) are much more probable than under a standard birth-death process. A corollary of this is that clade size a typical species ‘experiences’ (i.e. the expected clade size of a randomly selected species) is c. 8 times greater than the mean clade size. In Figure 1 we indicate both the mean clade size and the mean experienced clade size for illustration. This result should be compared to the equivalent result from a standard BDP where the mean ‘experienced’ clade is only two times greater than the mean [28]. This implies that the large majority of species we might encounter and/or study are contained in extremely large clades. Since clades are hierarchically structured this also implies that the diversity of any clade is likely to be dominated by its largest sub-clade. To illustrate this we consider a clade composed of two sister-groups of the same age, and calculate the expected proportion of the total diversity that is contained in one sister-group chosen at random. As shown in Figure 1B, the probability that a given proportion of total diversity is contained in a given sister-group is peaked strongly close to zero and one, indicating that one sister-group or the other typically contains the large majority of species in the clade as a whole. For example, there is a c. 50% chance that the larger sister group is at least 20 times larger than the other. This can be compared to the equivalent result under a standard BDP, in which the proportion of diversity contained in one sister-group is uniformly distributed between zero and one, and thus the probability of such an imbalance is only 10%. This implies that diversity among clades of the same age tends to follow the Single Big Jump principle [33], whereby sums of heavy-tailed random variables are dominated by their largest component.

**Figure 1:**
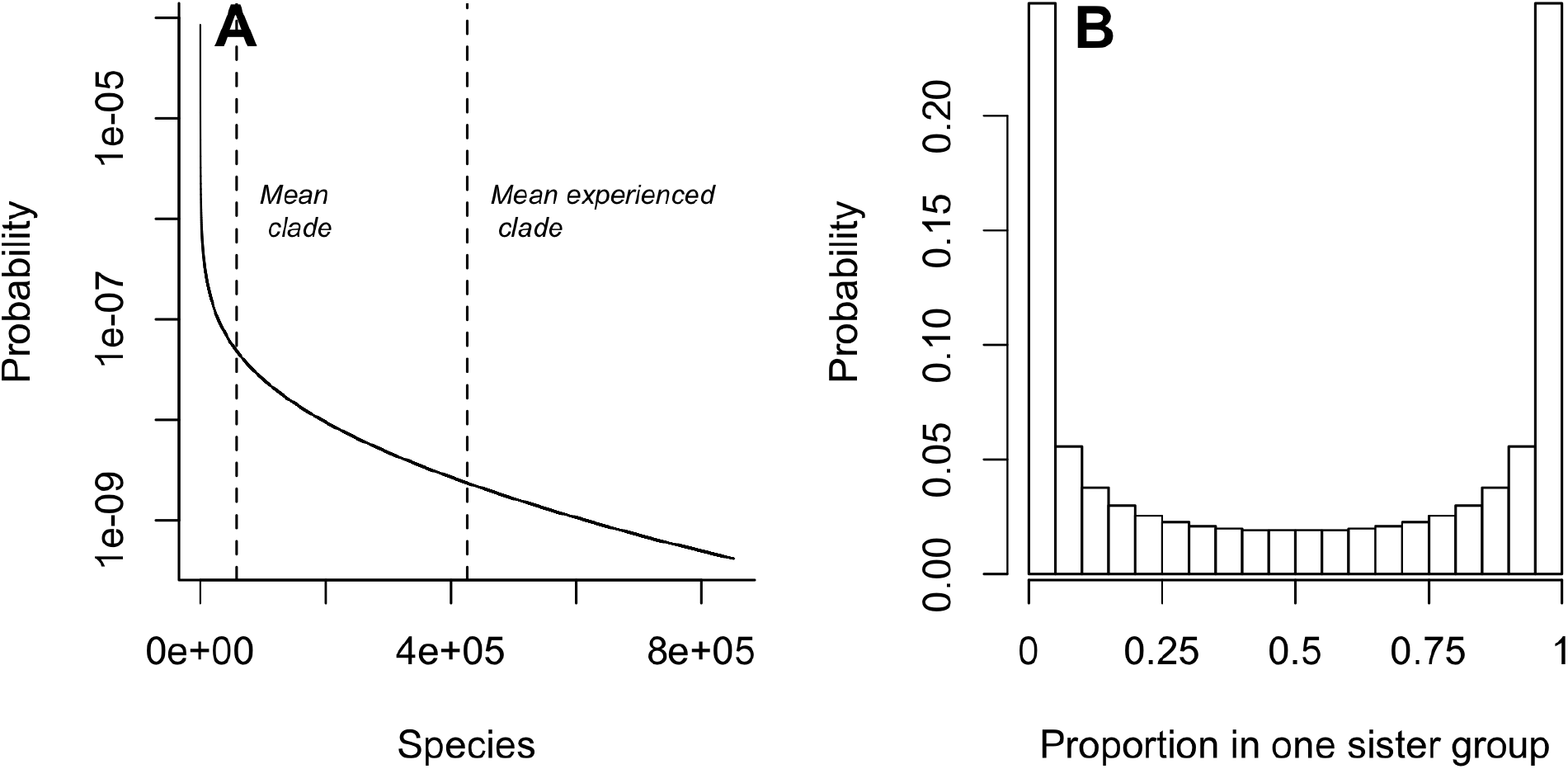
(A) Distribution of the number of species generated in clades that survive 500 myrs, with parameters *λ* = 0.51, *µ* = 0.5, *θ* = 0.01, *s* = 1, and an initial log-tempo *x* = 0. Note the log scale on the y-axis. The distribution is long-tailed and is characterised by a high probability of few species (*P* (*n <* 1000) ≃ 1*/*3) and a long tail allowing some very large clades to be generated (*P* (*n >* 50, 000) ≃ 1*/*4). The dashed lines indicate the mean clade size (c. 60,000) and the mean experienced clade size of a randomly chosen taxon (c. 400,000), indicating that most taxa are found in very large clades. (B) The probability distribution for the proportion of diversity contained within one randomly chosen sister group of a crown group, indicating that clades are typically highly imbalanced, with one sister group being much larger than the other.

### Diversification through time

The above analysis reveals the expected pattern of diversity in clades of a fixed age (500 myrs) which all start from a common ancestor with a typical tempo (*x* = 0). How does this pattern change through time, and between clades with different initial tempos? To explore these questions we focused on how the expected clade size varies through time for different initial values of *x*. We numerically solved equation 4 to obtain the expected clade size as a function of time values 0 *< t <* 500 myrs, and for different initial values of *x*_0_ ∈ {−2, 0, 2 }. In Figure 2A we show how the mean clade size varies through time for different initial tempos including clades that have gone extinct before the time in question. In Figure 2B we show the variation in the mean number of species through time conditioned on knowing that the clade survives to the present day (solid lines), and also the expected number of *lineages* (dashed lines) through time– these are species that have at least one descendant in the present day, and form the ‘reconstructed process’ that can (in principle) be inferred from modern molecular data. Clades that survive to the present experience the Push of the Past [28], a initial period of increased diversification when the clade is small. These results show the initial tempo has a substantial impact on how the clade diversifies and its eventual expected size. As we would intuitively expect, clades with high tempos initially diversify more quickly, and conversely those with low tempos diversify slowly. However, after some period of time the rate of diversification becomes stable; initially high tempo clades slow down and initially low tempo clades speed up, such that all clades eventually diversify at the same fixed rate, as seen in emergence of parallel lines of growth from all three initial conditions.

**Figure 2:**
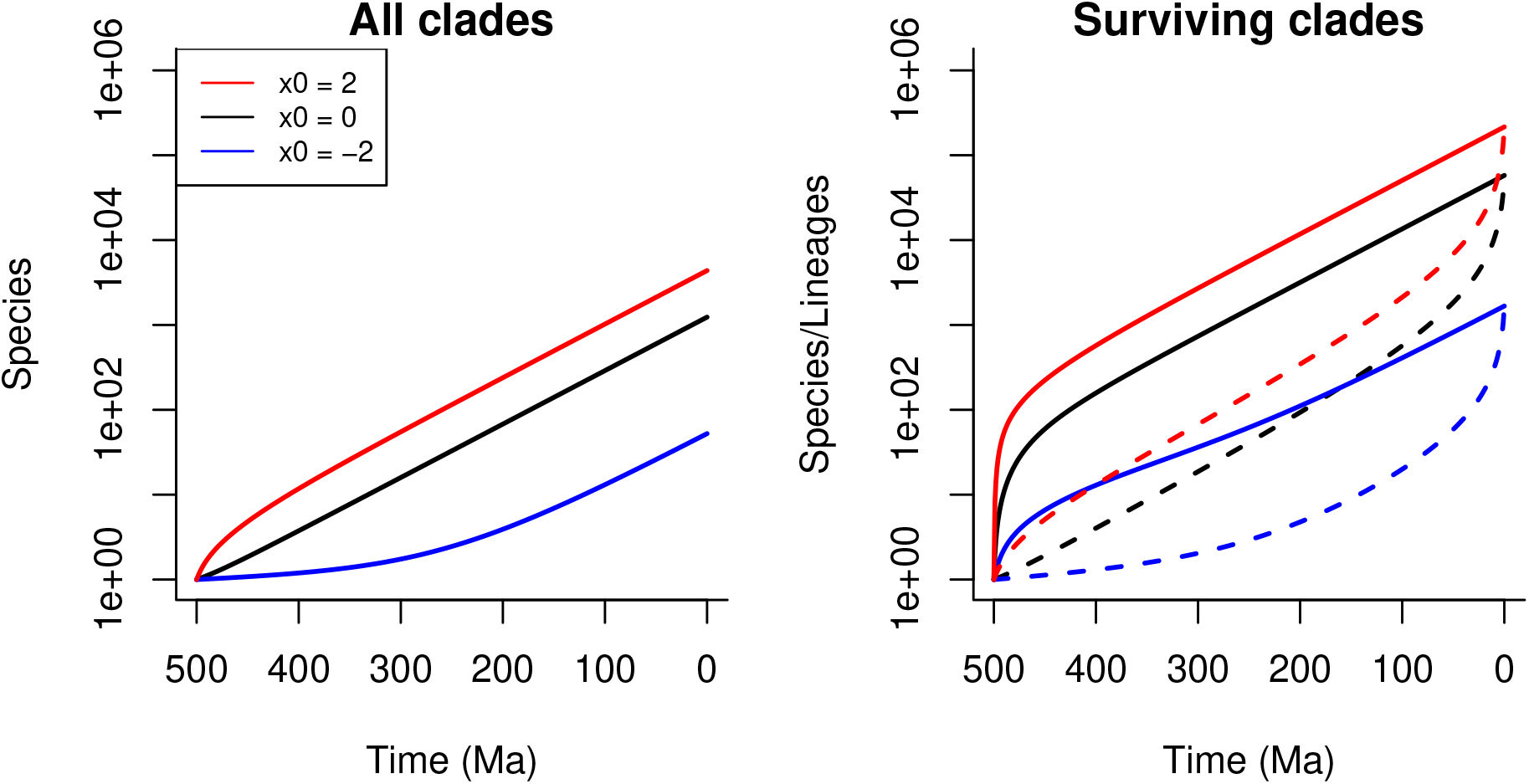
Diversification through time as a function of starting tempo. (A) The expected number of species through time for *x*_0_ =− 2 a clade starting 500 Ma with different initial log-tempos: *x*_0_ = 2 (blue line); *x*_0_ = 0 (black line); *x*_0_ = −2 (red line). These expectations include clades that are extinct. Clades with a higher starting tempo initially diversify more quickly (on average); eventually diversification stabilises to a fixed rate independent of the starting tempo. (B) Expected diversification profiles for clades that survive to the present day. Solid lines indicate the expected number of species through time; dashed lines indicate the expected number of lineages – species with surviving descendants. Surviving clades of all starting tempos experience the Push of the Past, mirrored by the Pull of the Present in the lineages. This effect is especially pronounced in the clades starting with the highest tempo.

The tempo of the root node of a clade therefore has transient effects that eventually decay as new species emerge whose own tempos diffuse away from the initial state. The duration of these transient effects is longer in clades that start with low tempos, since all processes including those that control the diffusion of tempos over time run slower. Although the effect of initial tempo is transient, it leaves an important signature in the eventual size of clades over the long term; because initially high tempo clades diversify more quickly in their early history they reach a larger size before reverting to a constant diversification rate, meaning that they have a much greater expected diversity in the present. This intuitively suggests that the largest clades of a given age in the present are likely to be those that originated from a high tempo common ancestor.

### Distribution of tempos over time

As a clade diversifies, the various taxa will develop different tempos as they diverge independently from the initial starting tempo, leading to a time-dependent distribution of log-tempos *p*(*t, x*). In the Methods we show that the evolution of this distribution obeys a replicator-mutation equation:

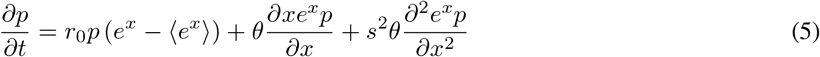

where the term 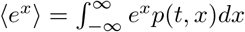 indicates the average value of *e*^*x*^ at a given time.

We numerically integrated this equation this probability through time 0 *< t <* 500 myrs for three initial starting log-tempos: *x*_0_∈ {−2, 0, 2} specified by initial conditions of the form *p*(*t* = 0, *x*) = *δ*(*x*−*x*_0_), where *δ*(·) is the Dirac delta function. The resulting evolution of the log-tempo probability distributions is shown in Figure 3. These results show that regardless of the starting tempo of the process, our model converges over time to the same stable distribution of log-tempos that is approximately normally distributed. Using the core set of model parameters described earlier gives a mean log-tempo of zero. When the process is initiated with a high tempo (*x* = 2) the convergence to this stable distribution is very rapid (red line). This is because the initially high tempo forces all processes to run fast, so time is effectively compressed. Conversely when the process is initiated with a slow tempo *x* = −2, the convergence is much slower, although the same stable distribution is eventually reached. This predicts the existence of long-lived substructures of the evolutionary tree in which evolution is effectively ‘running slow’. If other evolutionary processes such as molecular and morphological change are also covariant to the tempo this would imply the emergence of ‘living fossils’ – lineages with minimal morphological or molecular change over very long periods of time. The fact that all processes eventually converge on a single stable distribution irrespective of their starting point suggests that we can consider any taxon selected at random as having a tempo drawn from this distribution.

**Figure 3:**
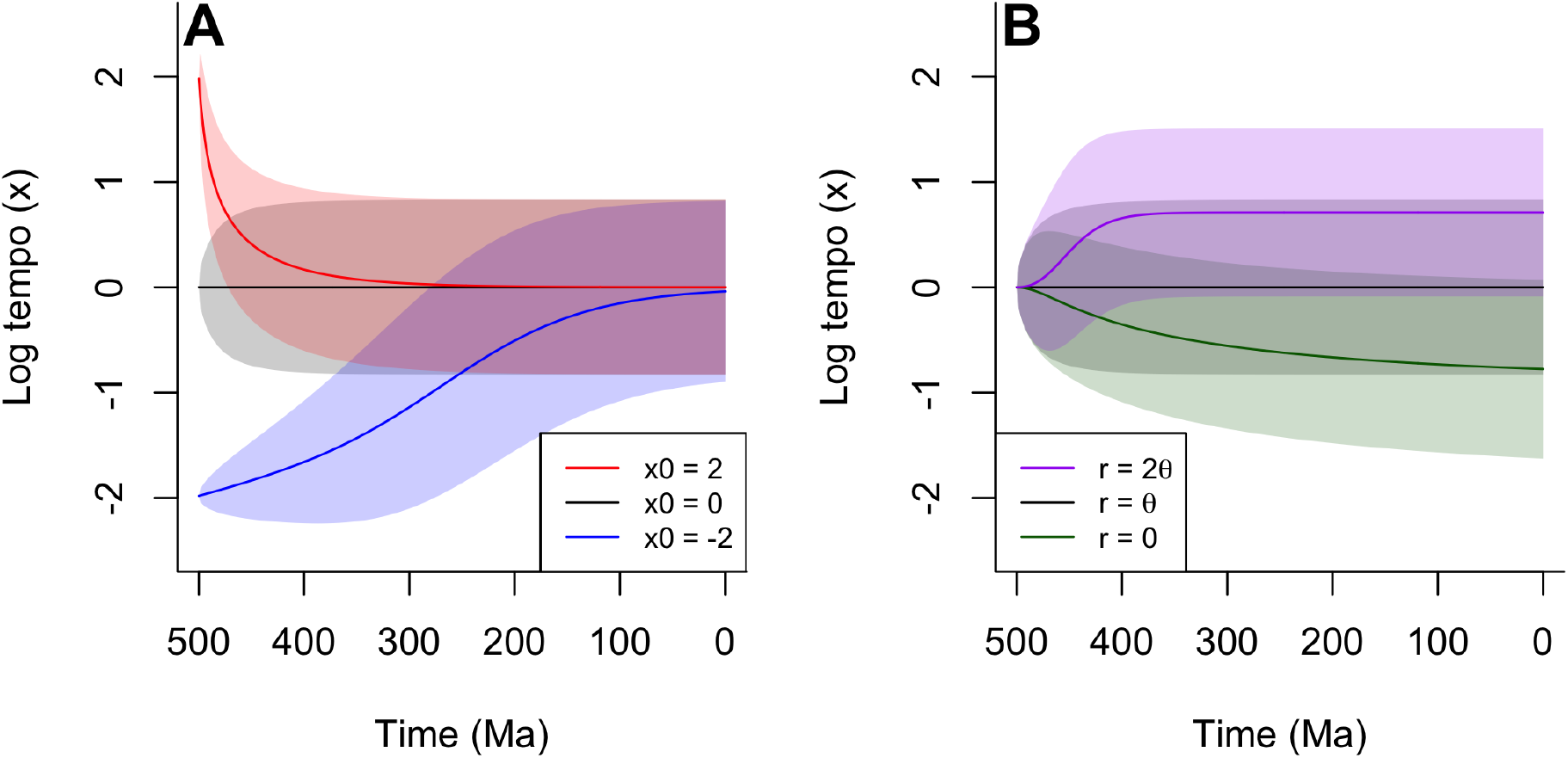
(A) The evolution of the distribution of log-tempos through time for clades starting from different initial log-empos: *x*_0_ =−2 (blue line); *x*_0_ = 0 (black line); *x*_0_ = 2 (red line). Lines indicate the expected log-tempo of a randomly chosen species, and shaded areas represent the standard deviation. Regardless of starting tempo, clades converge to the same equilibrium distribution of log-tempos. This convergence is fast in clades that start with high tempos. (B) Evolution of the log-tempo distribution for clades with different values of the diversification parameter *r*_0_, with a fixed value of *θ* = 0.01. Starting from the same tempo (*x*_0_ = 0), clades converge to different log-tempo distributions depending on the value of *r*_0_; higher values of *r*_0_ produce higher average tempos.

Varying the parameters of our model produces changes in the stable distribution of tempos. In particular the mean of this distribution increases with *r* and decreases with *θ* (Figure 3B); the limiting case where *r*_0_ = 0 the mean log-tempo can be shown to converge to -1 in closed form (see Methods, equation 5). The dynamics of diversification thus keep the mean tempo elevated relative to where it would be in the absence of diversification, since higher tempo lineages produce more descendants on average per unit time, which inherit the same high tempo from their parents nodes. An interesting corollary to this point is that without any sort of mean reversion process, tempos (and thus diversification rates) would simply tend to rapidly increase without limit. As this is not observed empirically, the suggestion must be that something tends to draw log-tempos towards a characteristic mean value (c.f. [10][2][34]). in the Methods we show that such a mean reversion can arise simply through entropic forces, without implying any necessary ecological mechanism.

### Patterns of historical tempo

So far we have considered what happens to various features of the evolutionary process as it is run forward from an particular initial condition. However, evolutionary analysis can be considered to be retrospective as well: one attempts to identify and explain patterns of evolution looking back in time from a vantage point in the present. As discussed by ref. [28] this perspective necessarily distorts the patterns we are likely to observe, especially if one also chooses to analyse clades that have unusual modern day properties. Such choices are common place: the most studied clades are often unusually diverse relative to clades of similar age; since most species are contained in these large clades they are often taken to be particularly representative of a particular epoch, despite in fact being high unrepresentative sample of clades in general.

To investigate the role that contingencies of clade selection have on the observed patters of evolution we considered two questions. First, if one randomly selects a species in the present and traces its lineage back in time, what expectations should we have about the evolutionary tempo of those ancestors? Second, what expectations should we have about the average tempo of earlier members of that clade overall? These are different questions since most historical taxa, even those with modern descendants (the lineages) will contribute little to modern diversity, owing to the Single Big Jump Principle [33] identified earlier (cf. Figure 1B). That is, the ancestors of most modern taxa constitute a very small subset of historical diversity.

Figure 4 illustrates our expectations about the historical patterns of tempo. Figure 4A shows the distribution of log-tempos for ancestors of a randomly chosen modern taxon. In the present these are centred around *x* = 0, which is the stable overall distribution of log-tempos shown in Figure 3A. As we look backwards in time the expected log-tempo of the ancestor rises sharply, before plateauing at *x* ≃0.6 at c. 100Ma. Conversely, the tempo of the clade as a whole tends to peak at its origin, as shown in Figure 4B. This illustrates the overall expected log-tempo of historical species within a clade inhabited by a typical modern taxon (i.e. one with a diversity equal to the mean experienced clade size). That is, the clades that contain most modern taxa are defined by a high early rate of evolution, which then undergoes a consistent secular decline to the present, while the direct ancestors of most modern taxa have uniformly elevated rates of evolution across the history of the clade until close to the present. A consequence of this result is that most modern taxa share relatively recent common ancestors (c. 100-150Ma), as they overwhelmingly tend to originate via a small subset of lineages that maintain high tempos until this point. This is despite the most recent common ancestor of *all* species being close to the origin of the clade.

**Figure 4:**
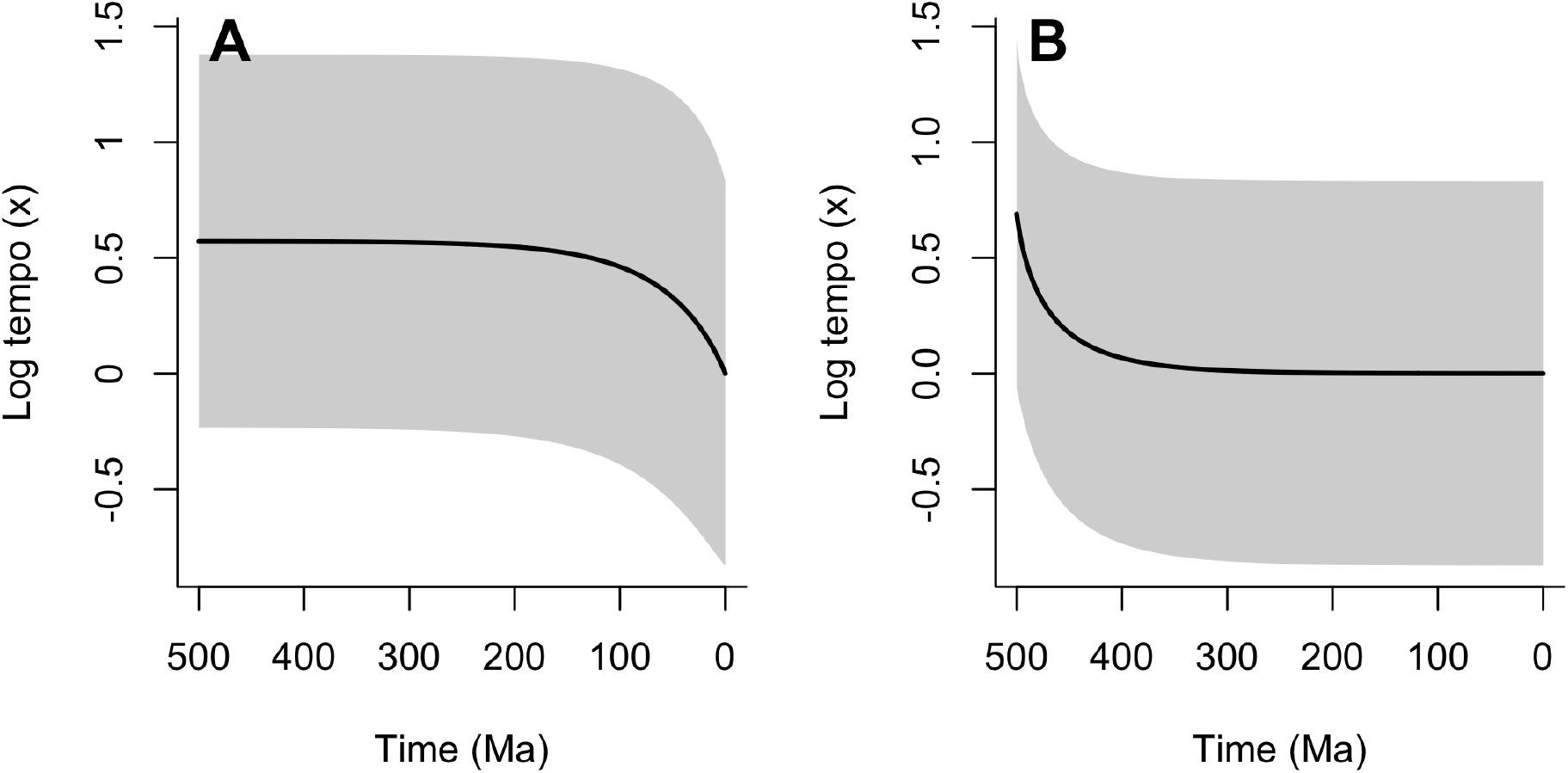
Expected patterns of historical tempo evolution, conditioned on present day observation choices. Lines indicate the expected log-tempo and shaded areas represent the standard deviation. (A) The expected historical log-tempo of ancestors of a randomly-chosen modern taxon, following its lineage back to the origin of the clade. Throughout this lineage expected log-tempos are elevated relative to the present day, declining rapidly shortly before the present. (B) Expected historical log-tempo of species in the clade as a whole. This is shown for a clade of the mean experienced clade size (the typical clade size of a randomly chosen modern species). Expected tempos are highest at the origin of the clade and decline through time as the clade diversifies.

### Effect of tempo variation on branch lengths and duration

So far we have considered the effect of tempo variation on the dynamics of the birth-death process, and by extension on diversification. We motivated our approach by noting that rates of molecular evolution are commonly assumed to vary in modern relaxed molecular clock analyses, and now we turn our attention to the interaction of molecular evolution and diversification. Specifically, we consider the expected duration (in real time) and amount of molecular change along branches with differing initial log-tempo values. In our model, tempo can vary within a branch, so the duration of branches is not necessarily exponentially distributed, in contrast to standard BDP models. Instead, the probability that a branch terminates (either by speciation or extinction) in a small interval of time Δ*t* depends on its current log-tempo and is given by *e*^*x*^(*λ*_0_ + *µ*_0_)Δ*t*.

As shown in Methods, this implies that the probability density *f*_*x*_(*t*) that a branch originating with log-tempo *x* terminates at time *t* obeys a partial differential equation of the form:

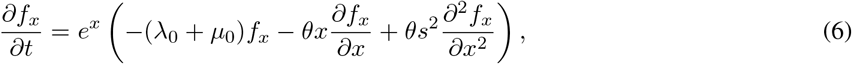

with initial condition *f*_*x*_(*t* = 0) = −*e*^*x*^(*λ*_0_ + *µ*_0_). In Figure 5A shows the solution to this equation for three different values of *x* ∈{−2, 0, 2}, illustrating the intuitive result that branches with lower initial tempos tend to have a greater duration – that is they exist for a longer time before either speciating or going extinct.

**Figure 5:**
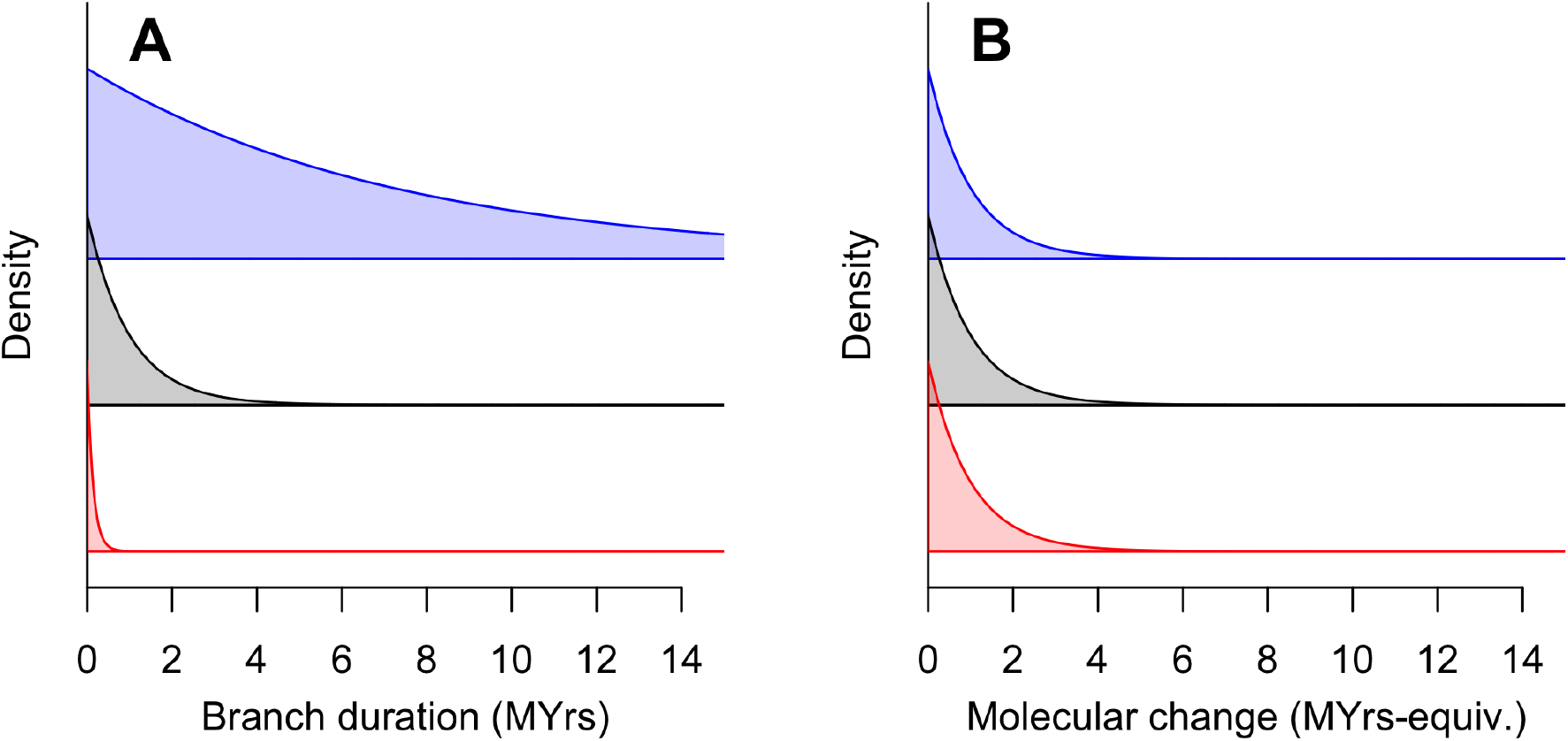
The distribution of branch durations (A) and amounts of molecular change along branches (B) for branches starting with different log-tempos: *x*_0_ =−2 (blue); *x*_0_ = 0 (black); *x*_0_ = 2 (red), assuming that molecular evolution is covariant with tempo. Branches that start with lower tempos are much longer on average in real time than those with high tempos. However, the expected amount of molecular change is independent of the starting tempo.

How does this effect of the initial tempo translate into the amount of molecular change that occurs within a branch? This is an important question, because the relationship between branch duration and molecular change is fundamental to the practice of molecular dating and more broadly to the inference of phylogenetic relationships based on the molecular genetic data from modern taxa [11].

If we assume that rates of molecular change co-vary with tempo alongside all other rates then the amount of molecular change Δ*w* that occurs in some small unit of time Δ*t* is given by:

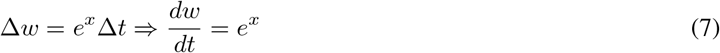

Applying a change of variables to express Equation 6 in terms of the molecular change *w* gives an equation obeyed by the probability density of molecular change *f*_*x*_(*w*) in a branch that starts with log-tempo *x*:

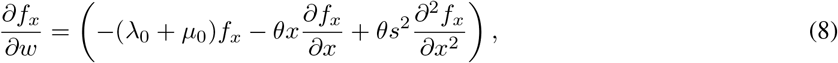

with initial condition *f*_*x*_(*w* = 0) = *λ*_0_ + *µ*_0_. Noticing that the partial derivatives in this equation will remain zero for all values of *w*, this simplifies to a standard exponential distribution:

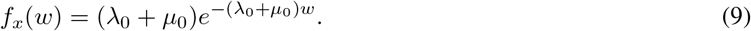

That is, the amount of molecular change contained in a branch is *independent* of the value of the tempo. This is illustrated in Figure 5B.

The key result then is that branches that start at higher tempos are typically shorter, but contain just as much molecular change, as longer branches that originate from lower tempos. This implies that a clade that starts with a high tempo is likely to be characterised in its early stages by short-duration branches that nonetheless contain just as much molecular change as later branches which are longer in duration. Since we have shown above that early high tempos are expected especially in clades that are particularly large, we can expect that this pattern to be commonly observed. As a corollary, if we further assume that morphological change also co-varies with tempo (c.f. [31, 21]) then the same pattern of rapid change along short early branches would be observed morphologically by an analogous argument.

## Discussion

We have described a model of macroevolution that allows the rates of speciation, extinction and molecular/morphological evolution to co-evolve through a variable evolutionary tempo parameter. This model provides a resolution to several outstanding difficulties in reconciling classic birth death models with empirical data. Allowing for tempo variation produces much greater variation in clade sizes over a given time horizon than under previous models, consistent with the fact that modern diversity is dominated by a relatively small number of very large clades across different taxonomic levels. An underappreciated consequence of this distribution is that if if we wish to understand how modern patterns of diversity arose, it is important to study the characteristic behaviour of such large clades, which, as we have shown here, differs markedly from that of clades as a whole. In other words, large and arguably charismatic clades such as arthropods, birds and angiosperms that are the subject of understandable interest, have quite different patterns of evolution than what an “average” clade might be inferred to have.

Our results show that these clades containing the bulk of modern diversity are likely to result from very high early evolutionary tempos, leading to short early branches (measured in real time). Because we conjecture that evolutionary tempo affects all rates in a covariant fashion, these short early branches are nonetheless expected to contain as much molecular and morphological change as later, longer branches, because the rates of molecular and morphological change are elevated in direct proportion to speciation and extinction. This offers an explanation for the observation of, for example, such elevated rates coupled in short early branches found in molecular studies that take the fossil record as a reliable guide to the age of the clade (e.g. [21]). We note that such studies tend to indicate a much older origin of the clade when the firm calibration based on the fossil record is removed; we hypothesise that this emerges because of the use of a model that assumes a homogeneous birth death process as the underlying description of diversification [26]. Modern molecular clock analyses typically employ a ‘relaxed-clock’ methodology that permits substantial changes in the rate of molecular evolution across time and between lineages, but these rates are decoupled from the rates of speciation, extinction and lineage creation (e..g [2]. Such a rigorous decoupling between evolutionary processes seems intuitively unrealistic, and indeed elevated rates of molecular evolution have been posited as a the *cause* of radiations [30], while in the fossil record morphological change is the key signature of diversification. As such, we argue that recognising the likely covariance between these rates is key to understanding apparent discrepancies between molecular signatures of diversification and the fossil record.

A covariant process that extends to rates of molecular evolution will produce similar amounts of molecular change on all branches of the tree, regardless of their duration in time. This suggests that from a molecular standpoint there will be little or no difference between a tree evolving according to homogeneous parameters over a longer period of time, or one that experiences rapid early evolution and diversification (or indeed an even older process that experienced very slow early evolution, although these will typically represent only a small proportion of modern diversity). As such, molecular data from modern taxa are unlikely to be able to discern which of these scenarios led to the molecular and species diversity we observe today. Precise and reliable fossil calibrations, in combination with molecular data can potentially reveal the typical distribution of rates within the time scope of those calibrations. However, extrapolation of younger rates into deeper time is problematic, as we have shown that these are likely to be higher in the past, beyond the deepest precise calibrations (c.f. [25]). This imposes a currently-insurmountable barrier to the use of the molecular clock for providing reliable clade age estimates, unless one can argue that rates of speciation and extinction are entirely decoupled from the more fundamental process of molecular change. As noted earlier, making such an argument would preclude many putative explanations for observed rapid radiations, as well as being counter-intuitive. We suggest therefore that the use of molecular clocks for making extrapolative deep-time age estimates is fundamentally unreliable (interpolations within a tree, between nodes of known age are likely to be more constrained, but here we expect that molecular data will add little to simpler methods).

As well as revealing the broad outlines of the dynamics of a varying tempo model of evolution, our analysis of this model also provides several empirical predictions:

1. Analysis of clades which are known to originate at similar times will show that the large majority of modern diversity is contained in a small subset of these clades. Most concretely, we anticipate that in pairs of sister groups, one group is likely to greatly dominate the diversity of the total (cf [35]).
2. The smaller sister group in a clade will be that which also experiences lower aggregate molecular and morphological change over its history. As such, the species on in this group will tend to retain more plesiomorphic features relative to those in the larger sister group. Potential examples of such a phenomenon include the onychophorans relative to arthropods, cyclostomes relative to gnathostomes [36], or priapulids relative to other ecdysozoans (e.g. [37]). This prediction gives some succour to the popular notion of “living fossils” that are slow evolving, have few species, and which to some extent resemble ancestral taxa (c.f. [38] for an alternative view).
3. The direct ancestors of most modern species will show elevated rates of evolution (diversification, molecular and morphological) throughout their history. Those historical species that have the most modern descendants will therefore show consistent rates of molecular evolution until close to the present, when they fall. However, if one analyses all historical taxa in a large clade (which is where most modern taxa reside) we expect to see very high rates of molecular change concentrated at the origin of the clade, declining consistently to the present. Nevertheless, both of these expected patterns take place within a wider context in which rates of evolution remain consistent *overall* – that is, measured over all species in all clades at a given time.
4. If we further assume that rates of evolution are associated with body size and generation time (e.g.. high rates being linked to small bodies and short generation times), we expect that a randomly chosen modern species will have experienced an increase in body size and generation time in the recent past, having probably originated from ancestors with smaller body size and shorter generation time.

Each of these predictions already enjoys some degree of empirical support in the existing literature, as indicated above. However, further research is needed to test each systematically to the extent that these predictions could be judged to be successful or falsified.

In conclusion, our analysis suggests that rate shifts are a crucial factor in explaining both current patterns of relative diversity among clades and historical patterns of diversification and molecular evolution, while removing the need to associate these rate shifts with special events or moments in time. By introducing a direct link between rates of molecular evolution and diversification, it unifies two key areas of statistical modelling within a common framework and points towards necessary developments in phylogenetic reconstruction and molecular dating in which this link is made explicit.

## Methods

### Generating functions

We will make extensive use of probability generating functions. A quick review of their important properties follows. A probability generating function, *G*(*z*) for the random variable *X* is defined as:

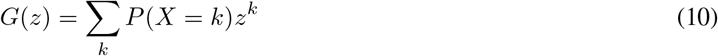

The probability generating function has several important properties that will be useful in the subsequent exposition. In particular:

i. Normalisation: *G*(*z* = 1) = Σ_*k*_ *P* (*X* = *k*) = 1 (in cases where *P* (*X* = *k*) represents a full probability distribution)
ii. Extinction probability: *G*(*z* = 0) = *P* (*X* = 0)
iii. Expectation: 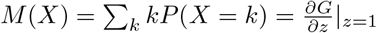
iv. Sum of random variables: If *W* = *X* + *Y*, then *G*_*W*_ (*z*) = *G*_*X*_ (*z*)*G*_*Y*_ (*z*)
v. Retrieval of probabilities: 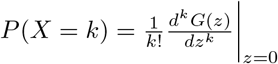

In respect of point (v) above, the values of *P* (*X* = *k*) can be retrieved efficiently by Fourier inversion:

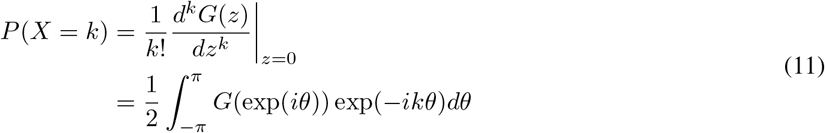

Where the integral expression makes use of the Cauchy integral formula. This expression can be efficiently solved numerically using Fast Fourier Transform methods [39]

### Derivation of equation specifying evolution of the generating function

Define *G*_*x*_(*t, z*) = Σ_*n*_ *P*_*n*_(*t, x*)*z*^*n*^ as the generating function for the number of species alive at time *t* from a process that starts at log-tempo *x* at time *t* = 0. We indicate the *x* dependence by means of a subscript for reasons of notational clarity in later analysis.

Assume that we know the generating function for all *x* at some time *t*. How will the generating function change over a small increment of time Δ*t*? Since the process is fundamentally homogeneous in time (i.e., there are no ‘special’ times’), we can construct this by considering a process that starts incrementally earlier than the known generating function. Within this small interval of time the process will change log-tempo incrementally according to an OU process, and furthermore may either speciate (producing two new independent processes with identical starting tempos) or go extinct. Given a current tempo *x*, the probability of speciation is *e*^*x*^Δ*tλ*_0_, and that of extinction is *e*^*x*^Δ*tµ*_0_. Based on these possible events, the new generating function is given by a mixture of generating functions at time *t*:

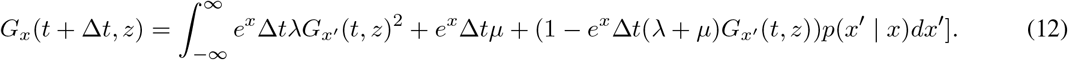

Here *p*(*x*^*′*^| *x*) specifies the probability for the tempo to transition from *x* to *x*^*′*^ over the time interval Δ*t*. We take *x* to evolve via an OU process, with correlation parameter *θ* and a stationary variance *s*^2^, experiencing an effective time *e*^*x*^Δ*t* within real time Δ*t*. Given this specification we have:

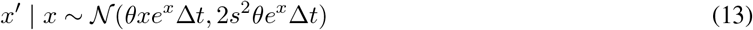

which yields: 𝔼[*x*^*′*^ − *x*] = −*θxe*^*x*^Δ*t* and 𝔼[(*x*^*′*^ − *x*)^2^] = 2*θs*^2^*e*^*x*^Δ_*t*_ up to first order terms in Δ*t*.

Taking a 2nd-order Taylor expansion of *G*_*x*_*′* (*t, z*)) around *x* and retaining first-order terms in Δ*t* gives:

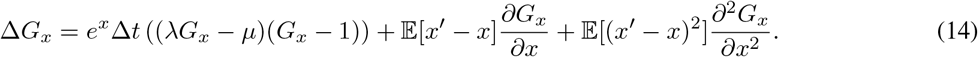

Where we have dropped the explicit dependence of *G*_*x*_ on arguments *t* and *z* for concision. Substituting the above expressions for 𝔼[*x*^*′*^− *x*] and 𝔼[(*x*^*′*^− *x*)^2^] and taking the limit as Δ*t*→ 0 gives the fundamental PDE of diversity evolution as given in equation 2.

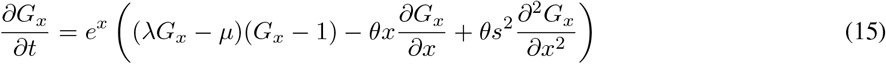

### Initial and boundary conditions

The most obvious question one can ask of this equation is: what is the probability that a process starting at log-tempo *x* will generate *n* species over time *t*? To answer this question we must solve equation 2 for different values of *z*, and use the Fourier inversion formula to retrieve the probability distribution *P*_*n*_(*x, t*). Solving equation 2 requires both initial and boundary conditions. For the question posed above the appropriate initial condition is given by *G*(*t* = 0, *x, z*) = *z* ∀*x*, since a process that does not evolve for any time must have one species. Choosing appropriate boundary conditions is more difficult; since we must solve equation 2 numerically we take ’no flow’ boundary conditions 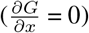 at some finite bounds *x*_min_ and *x*_max_ (we will usually use −10 *< x <* 10).

We can also ask how many species of log-tempo *y* will be produced at time *t* by a process that starts with log-tempo *x* at time *t* = 0. Define the generating function of this distribution by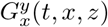. Some consideration will show that the time evolution of 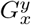 obeys the same PDE as that of *G*_*x*_, but with a different initial condition. Since a process that starts with log-tempo *x* cannot instantaneously evolve to one of *y* ≠*x*, we use the initial condition: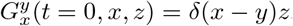, where *δ*(·) is the Dirac delta function.

### Evolution of the mean diversity

The mean of a distribution is straightforwardly recovered from its generating function via the relationship 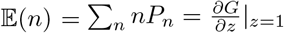. Applying this to the equation derived above for the evolution of the generating function gives the evolution of the mean diversity for a process that starts with log-tempo *x*. Defining *N*_*x*_(*t*) ≡𝔼(*n*| *x, t*) as the expected value of *n* at time *t* for a process starting with log-tempo *x*:

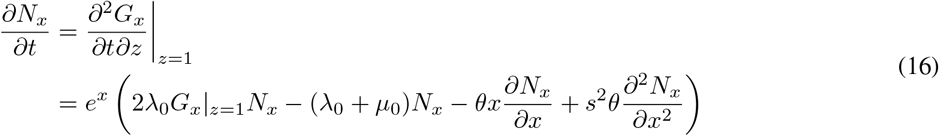

Since *G*|_*z*=1_ = 1∀ *x, t* by definition, we can simplify this to the expression given in equation 4:

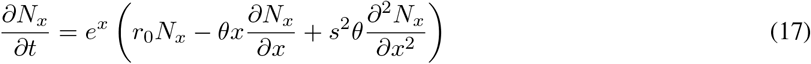

where *r*_0_ = *λ*_0_ − *µ*_0_.

By using initial conditions *N*_*x*_(*t* = 0) = 1∀ *x*, solving this equation gives the mean number of species generated by a process starting at time *t* = 0 and log-tempo *x*. As with the discussion of initial conditions above, we can also apply the same equation with different initial conditions to consider how many species with specific log-tempo *y* are generated by a process that starts at log-tempo *x*. Denoting the expected number of such species of this type as 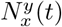 in this case we use the initial condition 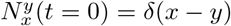, analogously to the case of solving for the generating function. By definition, the expected number of species in total will be the sum over all final log-tempos: 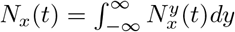. Furthermore, we can ask what the expected number of species with log-tempo *y* is at time *t* if the starting log-tempo is unknown but specified by a probability distribution *p*(*x*). In this case we have:

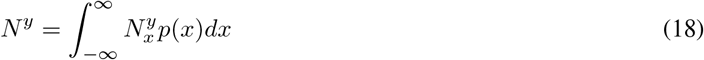

and the expected total number of species (considering all possible starting and current log-tempos) can be denoted simply as *N* (*t*) and is given by:

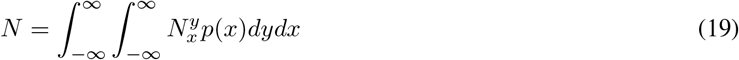

### Conditioning on survival

Equation 4 describes the evolution of the mean number of species through time, including all cases where the process goes extinct before the current time. If we want to ask how many species will be alive at time *t*, assuming that the process hasn’t gone extinct, we can do so straightforwardly by excluding the extinct cases:

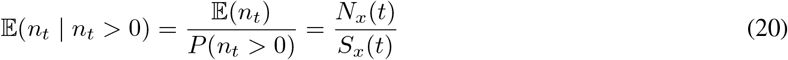

where *S*_*x*_(*t*) = 1 −*G*_*x*_(*t, z* = 0) is the survival probability for a process starting at log-tempo *x*, determined from solving equation 2 for *z* = 0. However, we may also want to know the expected number of species at some time *t*, conditioned on knowing that the process will survive to some future time *T*. In this case the conditioning is more complex. We make use of the identity:

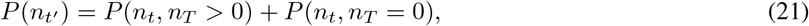

which leads to:

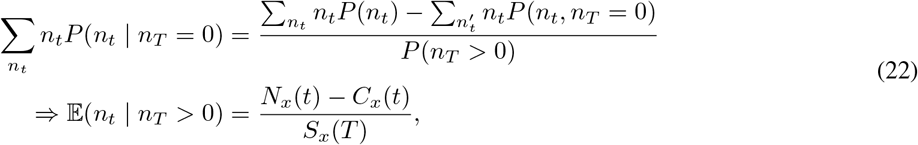

where *C*_*x*_(*t*) is a correction term depending on *x* and *t* that we need to determine. Define a new generating function 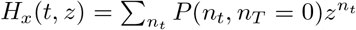. Differentiating *H*_*x*_ with respect to *z* and evaluating at *z* = 1 gives the required correction term in the equation above. As with the generating function *G*, the evolution of *H* is governed by equation 2:

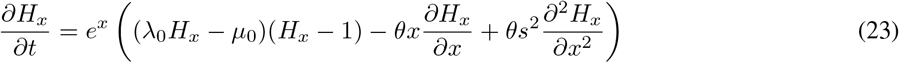

Differentiating with respect to *z* gives:

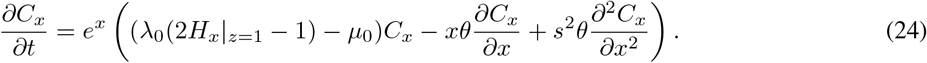

Unlike in the case for *G*_*x*_, *H*_*x*_| _*z*=1_ varies as a function of *x* and *t*, and so solution of this equation for *C*_*x*_ requires simultaneously solving this PDE and 2 with initial conditions: *H*_*x*_(*t, z*) = *zG*_*x*_(*T*− *t, z* = 0) and *C*_*x*_(*t*) = *G*_*x*_(*T* −*t, z* = 0).

### Lineages

Lineages are species in the past that have descendants in the present. Since molecular studies are based on extant species, any phylogeny reconstructed from these must consist of lineages. The evolution of lineages has thus been dubbed the ‘reconstructed process’ [8], since these constitute the phylogeny that can, in principle, be reconstructed from molecular or morphological analysis of modern taxa.

We are interested in the number of species alive at time *t* which will have descendants at some later time *T*. Recall 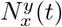 is the expected number of species of log-tempo *y* at time *t* in a process that starts at log-tempo *x*. The expected number of these that will have descendants at time *T* is *S*_*y*_(*T* −*t*) (the survival probability over time *T*− *t* for a new process starting with log-tempo *y*). Thus the expected number of lineages of log-tempo *y* at time *t* is 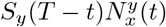. Summing over values of *y* gives the total expected number of lineages, *M*_*x*_(*t*) at time *t* for a process starting with log-tempo *x*, viewed from the perspective of time *T* (we leave this dependence on the time of observation implicit in the notation, but note that lineages are only defined from the perspective of a specific point in time):

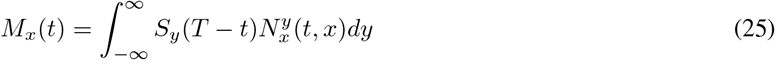

This expectation includes the cases where the number of lineages is zero, i.e where there are no species at time *T*. If we wish to condition on the process surviving to the present we must remove these cases by dividing by *P* (*n*_*T*_ *>* 0) = *S*_*x*_(*T*)

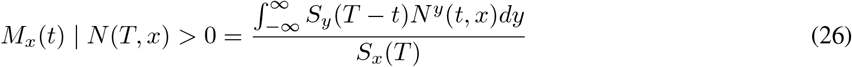

### Evolution of tempo distribution

Assuming that we start a process with log-tempo *x*, over time species generated by that process will diverge in tempos. How does this distribution of tempos evolve?

Consider starting a process with log-tempo *x*, and then selecting a species at random at some time *t*. The probability that this species has log-tempo *y* is given by:

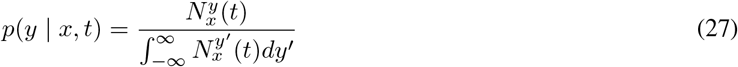

If the starting log-tempo is unknown, but drawn from a distribution *p*(*x*), then we can marginalise the above equation with respect to *x* to find the later distribution *p*(*y* | *t*):

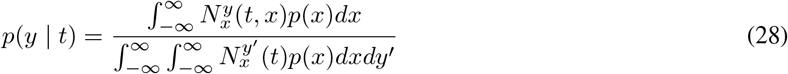

Taking the derivative with respect to time gives:

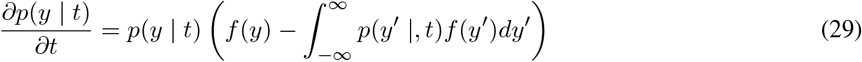

where 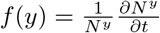. That is, the distribution of log-tempos evolves according to a replicator equation, where the ‘fitness’ of a log-tempo *y* is given by the proportional increase in 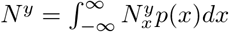.

If we assume that at some point in time the distribution of log-tempos is given by *p*(*x*), we can consider the instantaneous evolution of *N*^*y*^ from this time. Defining the current time to be *t* = 0, we have the initial condition:

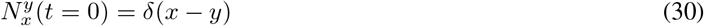

From equation 4, this implies that:

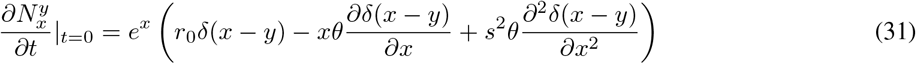

Applying standard rules for the operation of derivatives of the Dirac delta function, we can marginalise the above equation with respect to the initial distribution *p*(*x*) to give:

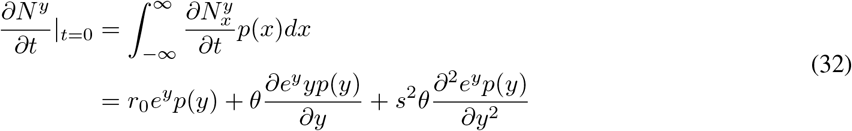

Substituting this into equation 29, and noting that again that 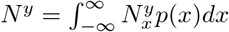, we get:

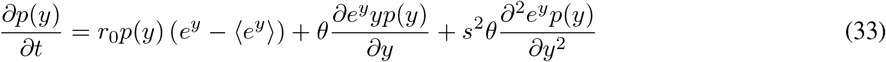

Where 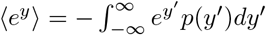 is the mean value of *e*^*y*^.

This then provides a replicator-mutation equation for the evolution of the tempo distribution, with the ‘fitness’ of log-tempo *y* being *r*_0_*e*^*y*^. In particular, it specifies that the stable long term distribution of log-tempos is given by the solution to:

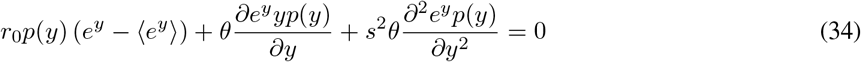

Notably, we can see that if *r*_0_ = 0, we recover the standard Fokker-Planck representation for the stationary OU process in the transformed distribution *e*^*y*^*p*(*y*):

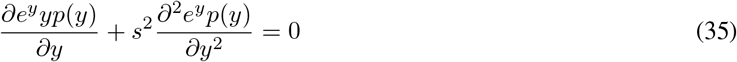

with the stationary solution 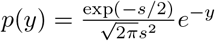 exp 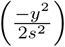 implying a mean log-tempo of ⟨*y*⟩ = −1. From the equilibrium equation we can also find another useful relationship on the mean value. Multiplying equation 34 by *y* and integrating gives:

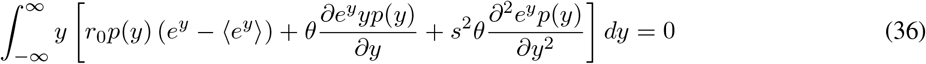

Integrating the partial differential terms by parts yields:

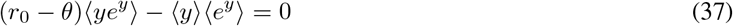

from which we can see that if *r*_0_ = *θ* then ⟨*y*⟩ = 0, i.e. the mean log-tempo will converge to zero when the diversification parameter is equal to the mean-reversion parameter.

### Inference of historical tempos

We assume that clades originate with an initial log-tempo *x*_0_ drawn from the stable distribution *p*(*x*) that we identified via analysis in the previous section (see Figure 3). This is equivalent to assuming that the common ancestor is a randomly chosen species resulting from the same process that has been running for sufficient time to reach an equilibrium distribution of tempos.

First we consider how likely it is that a species alive today at time *T* originated from an ancestor at time *t* with log-tempo *x*. Since we assumed that the clade originates with an ancestor drawn from the equilibrium distribution *p*(*x*), the prior probability that a species alive at time *t* has log-tempo *x* remains *p*(*x*) by definition of the equilibrium. We can determine a posterior estimate for the ancestor’s log-tempo by application of Bayes rule:

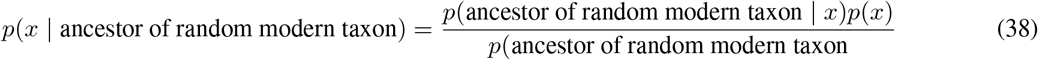

The likelihood term in this equation, *p*(ancestor of random modern taxon | *x*) is proportional to the expected number of modern species that an ancestor at time *t* will generate, *N*_*x*_(*T* −). This means we can rewrite the above as:

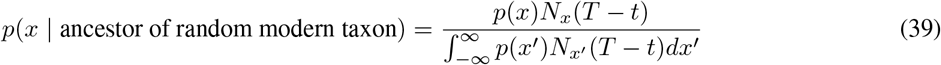

The equation above estimates the log-tempo of a direct ancestor of a modern taxon. We can also ask what the tempo of a randomly chosen member of the clade in the past is. This is a different question since only a small fraction of historical species will give rise to the majority of species alive today (see Figure 1). To estimate this we consider the probability of generating *n*_*T*_ species at time *T* from any starting log-tempo (based on solution of the generating function *G*_*x*_) and the probability that a randomly chosen species at time *t* has log-tempo *x*_*t*_ if the process starts at *x*_0_, *p*(*x*_*t*_| *x*_0_). From these probabilities we can infer the probability of a historical log-tempo *x*_*t*_ conditioned on the current diversity *n*_*T*_, using Bayes formula and marginalising over the unknown starting log-tempo *x*_0_:

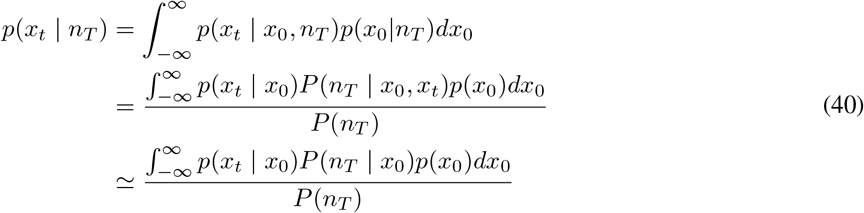

where the final approximation assumes that *P* (*n*_*T*_ | *x*_0_, *x*_*t*_) ≃ *P* (*n*_*T*_| *x*_0_). In general this approximation will be reasonable, because of the earlier result that modern diversity arises from a small subset of historical taxa. If the historical number of species at time *t* is high, a randomly chosen taxon is unlikely to contribute significantly to modern diversity and we can therefore treat *n*_*T*_ as being independent of this species and its tempo. Because of the Push of the Past [28], surviving clades will rapidly reach this state, and in the special case where *t* = 0 (i.e. the origin of the clade) the approximation holds exactly.

### Branch duration and expected molecular change

Considering a branch that begins with log-tempo *x*, what is the expected time until that branch terminates, either by speciation or extinction? For a branch to endure for time *t* + Δ*t* it must first fail to terminate in time Δ*t*, and then survive for a further time *t* with some new log-tempo *x*^*′*^. Integrating over the possible values of *x*^*′*^ we have:

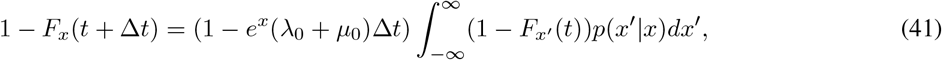

where *F* (*t* | *x*) is the cumulative probability that the branch originating with log-tempo *x* has terminated by time *t*. Taking a second-order Taylor expansion of *F*_*x*_*′* (*t*) around *x*^*′*^ = *x* and retaining first order terms in Δ*t* we have:

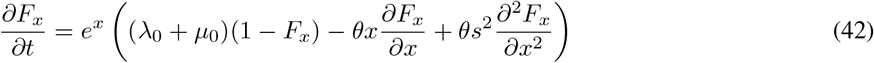

The probability density for the branch to terminate at time *t* is given by differentiation of the cumulative distribution: 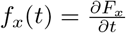. Applying this transformation to the equation above yields:

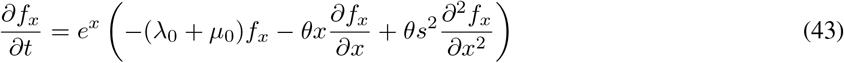

The probability density for a branch to terminate at time *t* thus follows the same form of differential equation as that for the mean number of species (equation 4), but with a negative ‘diversification rate’ of −(*λ*_0_ + *µ*_0_). Solving this equation requires the initial condition *F*_*x*_(*t* = 0) = 0 ∀*x*, which implies *f*_*x*_(*t* = 0) = *e*^*x*^(*λ*_0_ + *µ*_0_).

Assuming that molecular rates of change are covariant to tempo (*e*^*x*^), for every increment of time Δ*t* the expected amount of molecular change Δ*w* (in arbitrary units that we label as myrs-equivalent; 1 myrs-equivalent being the expected molecular change in 1 myrs at a fixed tempo of *τ* = 1) is Δ*w* = *e*^*x*^Δ*t*. We can transform the above equation for *F*_*x*_(*t*) (which is given in terms of real time *t*) into one that applies over *w* via a change of variables, to give the cumulative probability *F*_*x*_(*w*) that a branch terminate before accumulating *w* units of molecular change.

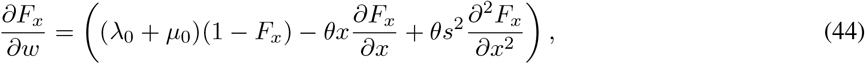

and as above we obtain the probability density to terminate at *w, f*_*x*_(*w*), by differentiation: 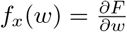:

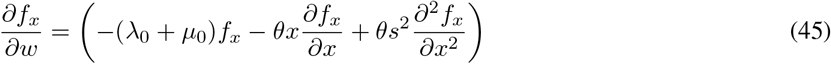

Here we have the initial condition *F*_*x*_(*w* = 0) = 0∀ *x*, which implies *f*_*x*_(*w* = 0) = *λ*_0_ + *µ*_0_. Consideration of this equation will show that the partial derivatives in *x* are initially zero and will remain zero for all values of *w*. Thus we can simplify the equation to:

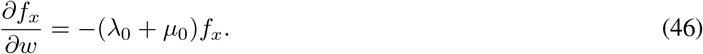

The solution to this equation is straightforward and shows that *w* follows an exponential distribution with rate *λ*_0_ + *µ*_0_:

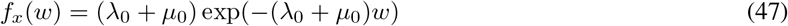

The notable feature of this density is that it does not depend on the starting log-tempo *x*, implying that the amount of molecular change in a branch is independent of tempo.

### Schematic for genetic encoding of tempo

Here we describe a simple model for how a genetic encoding of tempo can lead to the modified OU process we take as the basis for tempo evolution. Consider a binary string of *n* bases represented as ’1’ or ’0’, and define *ρ* as the proportion of bases that are ’active’ – that is, encoded as ’1’. We assume that these bases mutate independently and neutrally, and with a rate that is covariant to the tempo *e*^*x*^, such that the probability for each base to mutate in a small interval of time Δ*t* is *qe*^*x*^Δ*t*.

If the number of active bases at time *t* is given by *nρ*_*t*_, then in the interval of time Δ*t* the number of bases that mutate from ’1’ to ’0’ is binomially distributed as *B*(*qe*^*x*^Δ*t, nρ*_*t*_), and similarly the number mutating from ’0’ to ’1’ is binomially distributed as *B*(*qe*^*x*^Δ*t, n*(1− *ρ*_*t*_)). If we take *n* to be large and Δ*t* to be small these binomial distributions can be approximated by normal distributions, such that the number of mutations from ’1’ to ’0’ is normally distributed with mean *qe*^*x*^Δ*tnρ*_*t*_ and variance *qe*^*x*^Δ*t*(1 − *qe*^*x*^Δ*t*)*nρ*_*t*_, and the number of mutations from ’0’ to ’1’ is normally distributed with mean *qe*^*x*^Δ*tn*(1 − *ρ*_*t*_) and variance *qe*^*x*^Δ*t*(1 − *qe*^*x*^Δ*t*)*n*(1 − *ρ*_*t*_).

The change in the number of active bases is given by the number mutating from ’0’ to ’1’, minus the number mutating from ’1’ to ’0’. Given the results above, this change is also normally distributed. Taking the limit as Δ*t* becomes infinitesimal (denoted *dt*) and retaining only terms first order in *dt* we have:

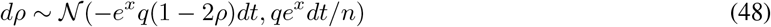

This is equivalent to the following form of stochastic differential equation:

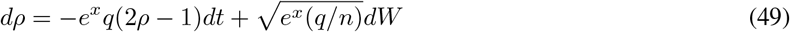

where *dW* is an increment from a standard Wiener process, with mean zero and variance *dt*.

To specify a genetic encoding of the log-tempo, let us now define *x* = *α*(2*ρ* − 1), where *α* is some arbitrary constant of proportionality, such that when half of bases are active this defines *x* = 0. We can then rewrite the above equation as:

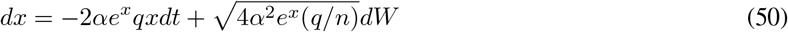

Defining new variables *θ* = 2*qα* and 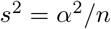, we have:

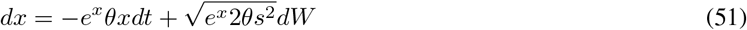

which is precisely the modified OU process specified in equation 1. By taking *α* to be sufficiently large we can extend the boundaries of minimum and maximum values of *x* such that arbitrarily high or low values of *x* are possible within this model. We have assumed that *n* is large, and this assumption means that boundary effects around *ρ* = 1 and *ρ* = 0 can be safely ignored as these states are highly unlikely to occur under a random mutation process.

This then provides a schematic representation of how tempo could be genetically encoded in a manner that naturally leads to the modified OU process description that we employ in this paper. The purpose of this schematic is not to argue that this represents the actual genetic encoding of tempo in any specific details, but instead to illustrate how such an encoding would naturally give rise to the mean-reversion properties of the OU process, via the action of entropic forces. That is, the log-tempo tends to revert to the mean not due to any ecological mechanism, but simply because there are more possible encodings with *x* ≃ 0 that those that encode more extreme values of *x*.

## Acknowledgments

This work was supported by Vetenskapsrådet (VR) grant 2022-03522 and UK Research and Innovation Future Leaders Fellowship MR/S032525/1 & MR/X036863/1. We are grateful to Tobias Uller, Ivan Prates, Dan Rabosky, Ben Slater and Jonathan Ward for discussions and help with literature for various aspects of this work. We are particularly grateful to Antonio Segalini, who was generous with his time and expertise in discussing and coding the numerical solution of PDEs.

